# Effect of Egg Irradiation on Development and Sterility of Wild-Type and *Wolbachia* Trans-Infected *Aedes aegypti* Mosquito Vectors

**DOI:** 10.1101/2024.09.23.614409

**Authors:** Pattamaporn Kittayapong, Suwannapa Ninphanomchai, Parinda Thayanukul, Limohpasmanee Wanitch

## Abstract

**Background:** Sterile Insect Technique (SIT), Incompatible Insect Technique (IIT) and a combination become the alternative promising vector control approaches. In these approaches, the targeted mosquitoes need to be sterilized and released. So far, the irradiation of mosquitoes has been conducted at the pupae or adult stages. In this study, we investigated the possibility of the X-ray irradiation at the egg stage and also assessed the effect on the development and sterility of both wild-type and *Wolbachia* trans-infected *Aedes aegypti* mosquito vectors.

**Methodology/Principal Findings:** The eggs of *Aedes aegypti*, both wild-type and *Wolbachia* trans-infected lines, were irradiated using X-ray at the doses of 1, 3, 5 and 7 Gy. Development of immature stages was observed. For wild-type *Aedes aegypti*, X-ray irradiation at the doses from 3 Gy decreased the development of the first instar larvae and increased the development of the third instar larvae but there was no effect on pupae when compared to those of the controls (*p* < 0.05). For *Wolbachia* trans-infected ones, the irradiation dose as low as 1 Gy could increase the development of the forth-instar larvae while the irradiation dose of 7 Gy induced significantly high mortality in pupae. To assess sterility, males and females emerged from irradiated eggs were mated with the normal ones. Results showed that the irradiation doses for sterilization of *Wolbachia* trans-infected *Ae. aegypti* was lower than those of the wild-type ones. The egg hatch rate of both males and females could significantly be reduced from the irradiation dose of 5 Gy in *Wolbachia* trans-infected *Ae. aegypti* and 7 Gy in the wild-type ones.

**Conclusions/Significance:** Our findings revealed the possibility to sterilize both wild-type and *Wolbachia* trans-infected *Ae. aegypti* by applying a low-dose X-ray irradiation at the egg stage. Egg irradiation could make an implementation of SIT, IIT or combined IIT/SIT for vector control much more feasible as the sterile dry eggs are easier to distribute and operate when compared to other developmental stages of mosquitoes.

**Author summary:** Releasing sterile males to control *Aedes aegypti* mosquitoes and the diseases they transmit is one of the alternative environmental friendly approaches. So far, sterilization of mosquitoes has been targeted on the pupae or adult stages. In this study, we investigated the possibility to apply X-ray irradiation at the egg stage which is more feasible and easier to operate. *Aedes aegypti* eggs, both infected and uninfected with the *Wolbachia* bacteria, were irradiated by X-ray at the doses of 1, 3, 5 and 7 Gy. Results showed that the irradiation dose of 7 Gy significantly caused more than 92% sterility in both wild-type males and females. However, this irradiation dose could be reduced to 5 Gy to sterilize both males and females infected with *Wolbachia*. In conclusion, the X-ray irradiation dose of 5 Gy could be an optimum dose for irradiation of *Wolbachia*-infected *Ae. aegypti* eggs as it could induce high level of sterility in males and nearly complete sterility in females while the irradiation dose of 7 Gy could induce sterility in both males and females uninfected with *Wolbachia*. This approach should be useful for the sterile male release programs to control *Aedes*-transmitted diseases.

## Introduction

*Aedes aegypti* is one of the most widespread and invasive mosquito species that is highly abundant in tropical countries [1, 2]. Significant public health important diseases, such as dengue, chikungunya and zika viruses, are transmitted by *Ae. aegypti* mosquitoes [3-7]. Currently, there were no highly efficient vaccines available for treatment of dengue, chikungunya, and zika viruses but several vaccine candidates are undergoing evaluation [8]. Regular immature surveillance and implementation of appropriate control measures have been recommended by WHO for vector control of *Aedes* mosquitoes [9]. However, challenges in insecticide resistance and treating small larval breeding sites have been highlighted [8, 10]. Over the past decade, several technologies have been developed to strengthen the conventional vector control methods [11], i.e., the release of fertile genetically modified mosquitoes (GMM) to alter the fitness of wild population [11, 12], the release of transgenic males to reduce the target populations [11, 13, 14], the exploit of cytoplasmic incompatibility induction properties of W*olbachia* by the incompatible insect technique (IIT) to enable the production of non-viable eggs [11, 15, 16], and the release of insects sterilized by radiation or the sterile insect technique (SIT) that rely on the release of sterile insects in order to reduce the reproduction of a natural population of the same species [11, 17]. Because of social concerns, cultural acceptance, and regulatory approval, the use of some methods (i.e., GMM, IIT) for vector control has encountered a number of challenge, therefore, the radiation-based approach was considered as the most feasible and safest solution [11].

An effect on radio sensitivity varies with age or age within the life stage [18] and different developmental stages are not equally affected by radiation [19]. However, due to a great variation in life cycle, life span, and exposure pathway, interaction between radiation and a wide range of biological species is not very well understood [20, 21]. On the other hand, production of sterile males for release through application of SIT approach has been focused and mainly obtained by irradiating pupae [22, 23, 24]. To date, only one study has been conducted to compare sensitivity of immature stages of *Ae. aegypti* with radiation, but this study was focused on gamma radiation [25]. The use of gamma radiation has become problematic because of government regulation, supply, usage and disposal of radioactive isotopes [24, 26]; as a result, an adequate alternative technology without the security risk such as X-ray technology has been urgently needed [26]. X-ray is among the ionizing radiation source that has been utilized in sterile insect releasing program [27] and X-ray based sterilization has been reported for a variety of insect pests including mosquitos [26, 28, 29, 30, 31]. However, only X-ray irradiation of pupae [28, 29, 30, 32] or adults [31] has been studied. To date, no X-ray studies have been conducted on eggs, pupae, or larvae of *Ae. aegypti* for comparison, as radiation has the effect on each developmental life cycle of *Ae. aegypti* [25], and impact on each life history trait of the species varies with dosage [20].

In this study, we aimed to investigate the effect of X-ray irradiation at the egg stage on development and sterility of *Wolbachia* trans-infected and uninfected *Aedes aegypti* mosquito vectors. Our findings will provide useful information on an optimal dose for radiation-induced sterility of *Ae. aegypti* at the egg stage which is the most appropriate and feasible for SIT, IIT or combined IIT/SIT implementation for vector control.

## Materials and methods

### Mosquito strain and rearing

The *Aedes aegypti* mosquitoes used in the present study were originally collected from several communities in Chatuchak District, Bangkok. A *Wolbachia* trans-infected *Ae. aegypti* colony was obtained from direct microinjection using the *Ae. aegypti* mosquito colony from Chatuchak District, Bangkok and the *w*AlbB *Wolbachia* strain from the *Ae. albopictus* colony originating from Plaeng Yao District, Chachoengsao Province, using the method described by Ruang-areerate and Kittayapong [30]. The establishment and characteristics of the *Wolbachia*-infected mosquitoes were demonstrated in Ruang-areerate and Kittayapong [33] and Kittayapong et al. [34].

Mosquitoes were reared in an aluminum mass-rearing cage sized 30 cm x 30 cm x 30 cm. in a screened climatic control insectary at the Center of Excellence for Vectors and Vector-Borne Diseases (CVVD), Faculty of Science, Mahidol University at Salaya, Nakhon Pathom, Thailand, with 75 ± 2% relative humidity, 27 ± 2°C, and a photoperiod of L12:D12, and were fed with 10% sucrose solution. Males and females were allowed to mate for 2-3 days; then the females were fed with pig blood by using the Hemotek membrane feeding system (Hemotek Ltd., UK) for 3-4 consecutive days after mating. The blood, obtained from a qualified slaughterhouse, was treated with 10% of EDTA (SCHARLAU, Spain) as an anticoagulant. Egg papers were placed in the containers inside the cage 1-3 days following blood-feeding. After 3-4 days, the egg papers were then collected, dried for 1-3 days at room temperature, and placed on a plastic tray prior to irradiation process.

### Irradiation and sex separation procedure

In this experiment, a total of 2,000 *Wolbachia* uninfected *Ae. aegypti* eggs, stored not more than one month, were counted and separated into 5 portions of 400 eggs. Four portions were irradiated with X-ray irradiator model RS 2400 (Rad Source Technologies, Inc. USA) at the irradiation doses of 1 Gy, 3 Gy, 5 Gy and 7 Gy respectively; and the remaining one portion was used as a control. Eggs were placed on a paper and then transferred into a plastic container volume 1,038.69 cm^3^ (diameter 11.5 cm) covered with a screened lid. Eggs were transported prior to irradiation by an air-condition car from the Center of Excellence for Vectors and Vector-Borne Diseases (CVVD), Faculty of Science, Mahidol University at Salaya, Nakhon Pathom Province to the Thailand Institute of Nuclear Technology (TINT) (Public Organization), Ministry of Higher Education, Science, Research and Innovation at Nakhon Nayok Province. The distance from CVVD, where the rearing facility was located, to TINT was about 100 km or 3 hours by car for a round-trip. Irradiation was conducted by the experienced and trained staff of TINT.

### Mosquito development process

After irradiation, eggs were transported back to CVVD. Then they were hatched in deionized water and newly emerged larvae were transferred into a plastic tray sized 32 cm x 42 cm x 5 cm with a total of 100 larvae per tray. Larval diets were provided with the amount between 0.5 -2.0 g per day according to the developmental stage. Dead larvae were removed by using a dropper and the number of dead larvae was counted and recorded daily. When larvae developed into pupae, male and female pupae were sex separated using the local pupal sex separator modified from the larval-pupal sex separator (John Hock Co., Ltd., USA). Then they were separately transferred into a mosquito cage sized 20 cm x 20 cm x 20 cm. The number of non-emerged male and female pupae was counted and recorded. After pupae became adults, 10% of sucrose solution was provided inside the mosquito cage.

### Sterility test

After emergence, *Wolbachia* uninfected *Ae. aegypti* aged 2-3 days were cross-mated according to the following mating pairs: 1) males emerged from irradiated eggs and non-irradiated females (IR Non-WolB M x Non-IR Non-WolB F), 2) females emerged from irradiated eggs and non-irradiated males (IR Non-WolB F x Non-IR Non-WolB M); and 3) non-irradiated males and non-irradiated females (Non-IR Non-WolB M x Non-IR Non-WolB F). Mosquitoes were allowed to mate for 2-3 days and then blood feeding was provided. Blood-fed females were individually transferred into a small plastic cup and the condition for oviposition was provided. Females were allowed to lay eggs for 3-5 days and the number of eggs laid per female was counted and recorded. Eggs were hatched as previously described and the number of hatched eggs was counted and recorded for assessment of sterility. Four replicates were conducted for each experiment.

The same experiments were conducted for those of *Wolbachia* trans-infected *Ae. aegypti* with three cross-mating pairs as follows: 1) males emerged from irradiated eggs and non-irradiated females (IR WolB M x Non-IR WolB F), 2) females emerged from irradiated eggs and non-irradiated males (IR WolB F x Non-IR WolB M), and 3) non-irradiated males and non-irradiated females (Non-IR WolB M x Non-IR WolB F).

### Statistical analysis

Data was entered and cleaned using Microsoft Office Excel 2016 and statistical analysis was performed further using SPSS 18.0 (Mahidol University License (Chicago, SPSS Inc.). Numbers of larvae, pupae, eggs, and egg hatch rate were analyzed by using paired-sample t-test; and Induce sterility (IS) was analyzed by using one sample t-test. P-values of less or equal to 0.05 were considered significant.

## Results

### Development of wild-type *Aedes aegypti* after being irradiated at the egg stage

Results showed that X-ray irradiation of the wild-type *Wolbachia* uninfected *Ae. aegypti* eggs with the irradiation doses from 3 Gy to 5 Gy significantly decreased the development of the first instar larvae, whereas the irradiation doses from 3 Gy to 7 Gy significantly increased the development of the third instar larvae when compared to those of the controls (Fig. 1, Table 1). When larvae developed into pupae, late development or high mortality of pupae was observed at the irradiation dose of 7 Gy when compared to other irradiation doses or the controls; but this difference was not statistically significant (data not shown).

**Figure 1.**
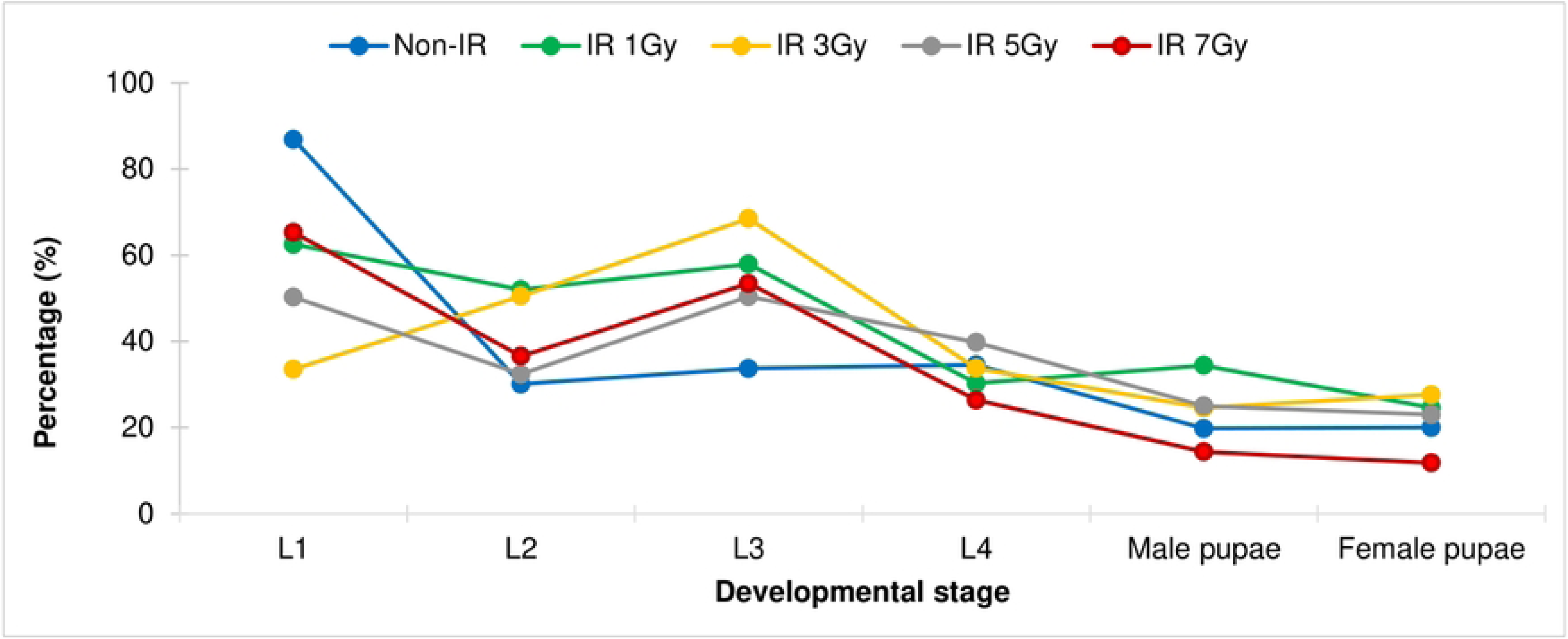
Comparison of larval and pupal development of the wild-type *Wolbachia* uninfected *Aedes aegypti* after eggs being irradiated with X-ray at the irradiation doses of 1 Gy, 3 Gy, 5 Gy and 7 Gy.

**Table 1.**
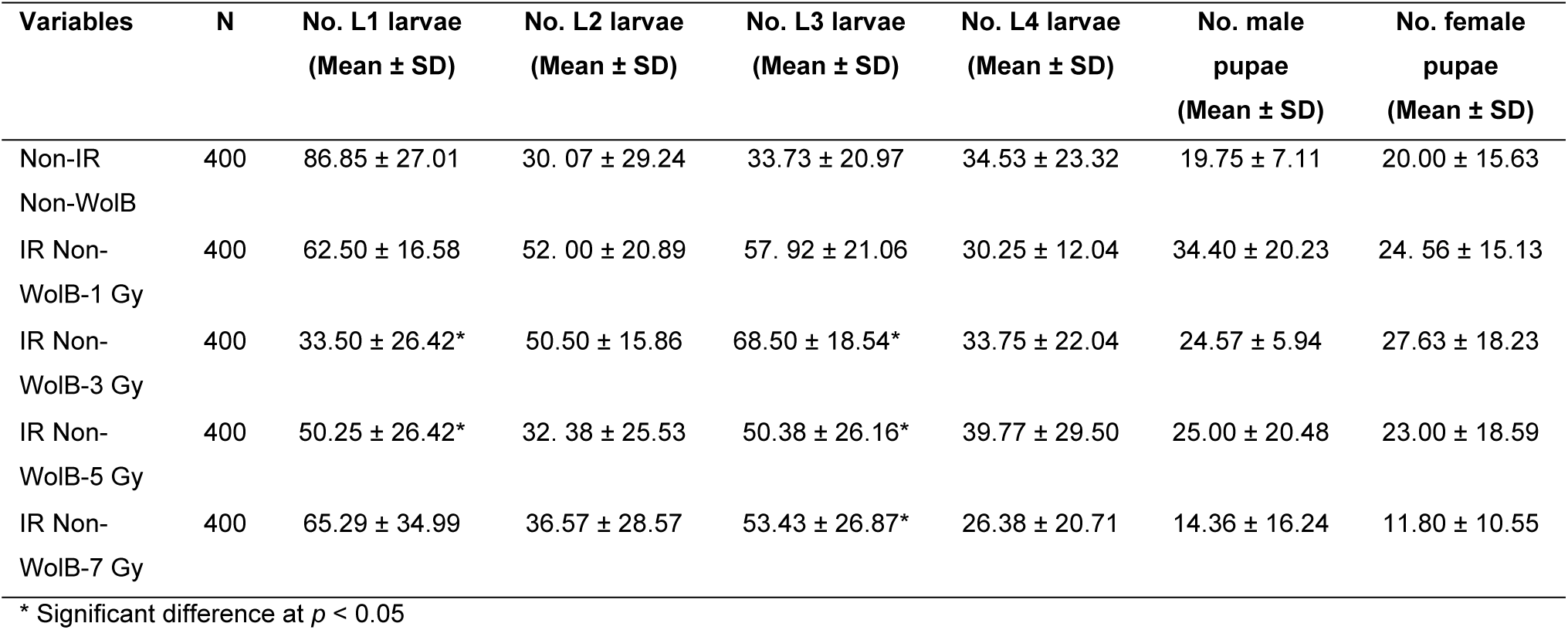
Comparison of larval and pupal development of the wild-type *Wolbachia* uninfected *Aedes aegypti* after eggs being irradiated with X-ray at the irradiation doses of 1 Gy, 3 Gy, 5 Gy and 7 Gy.

| Variables | N | No. L1 larvae<br>(Mean ± SD) | No. L2 larvae<br>(Mean ± SD) | No. L3 larvae<br>(Mean ± SD) | No. L4 larvae<br>(Mean ± SD) | No. male pupae<br>(Mean ± SD) | No. female pupae<br>(Mean ± SD) |
| --- | --- | --- | --- | --- | --- | --- | --- |
| Non-IR Non-WolB | 400 | 86.85 ± 27.01 | 30.07 ± 29.24 | 33.73 ± 20.97 | 34.53 ± 23.32 | 19.75 ± 7.11 | 20.00 ± 15.63 |
| IR Non-WolB-1 Gy | 400 | 62.50 ± 16.58 | 52.00 ± 20.89 | 57.92 ± 21.06 | 30.25 ± 12.04 | 34.40 ± 20.23 | 24.56 ± 15.13 |
| IR Non-WolB-3 Gy | 400 | 33.50 ± 26.42* | 50.50 ± 15.86 | 68.50 ± 18.54* | 33.75 ± 22.04 | 24.57 ± 5.94 | 27.63 ± 18.23 |
| IR Non-WolB-5 Gy | 400 | 50.25 ± 26.42* | 32.38 ± 25.53 | 50.38 ± 26.16* | 39.77 ± 29.50 | 25.00 ± 20.48 | 23.00 ± 18.59 |
| IR Non-WolB-7 Gy | 400 | 65.29 ± 34.99 | 36.57 ± 28.57 | 53.43 ± 26.87* | 26.38 ± 20.71 | 14.36 ± 16.24 | 11.80 ± 10.55 |

\* Significant difference at $p < 0.05$

### Sterility of wild-type *Aedes aegypti* after being irradiated at the egg stage

In terms of sterility of males emerged from irradiated eggs, it was found that the irradiation doses starting from 3 Gy onward significantly reduced the total number of eggs and the total number of hatched eggs of the cross-mating pairs between males emerged from irradiated eggs and non-irradiated females (IR Non-WolB M x Non-IR Non-WolB F), with a significant reduction of egg hatch rates at the irradiation doses starting from 1 Gy onward (0.59 ± 0.14 vs 0.70 ± 0.09, df = 29, t = 3.808, *p*-value = 0.001) (Fig. 2, Table 2). However, males became nearly complete sterile at the irradiation dose of 7 Gy (IS = 99.92; 0.08 ± 0.18 vs 0.70 ± 0.09, df = 16.693, t = 29, *p*-value = 0.000) (Fig. 2, Table 2).

**Figure 2.**
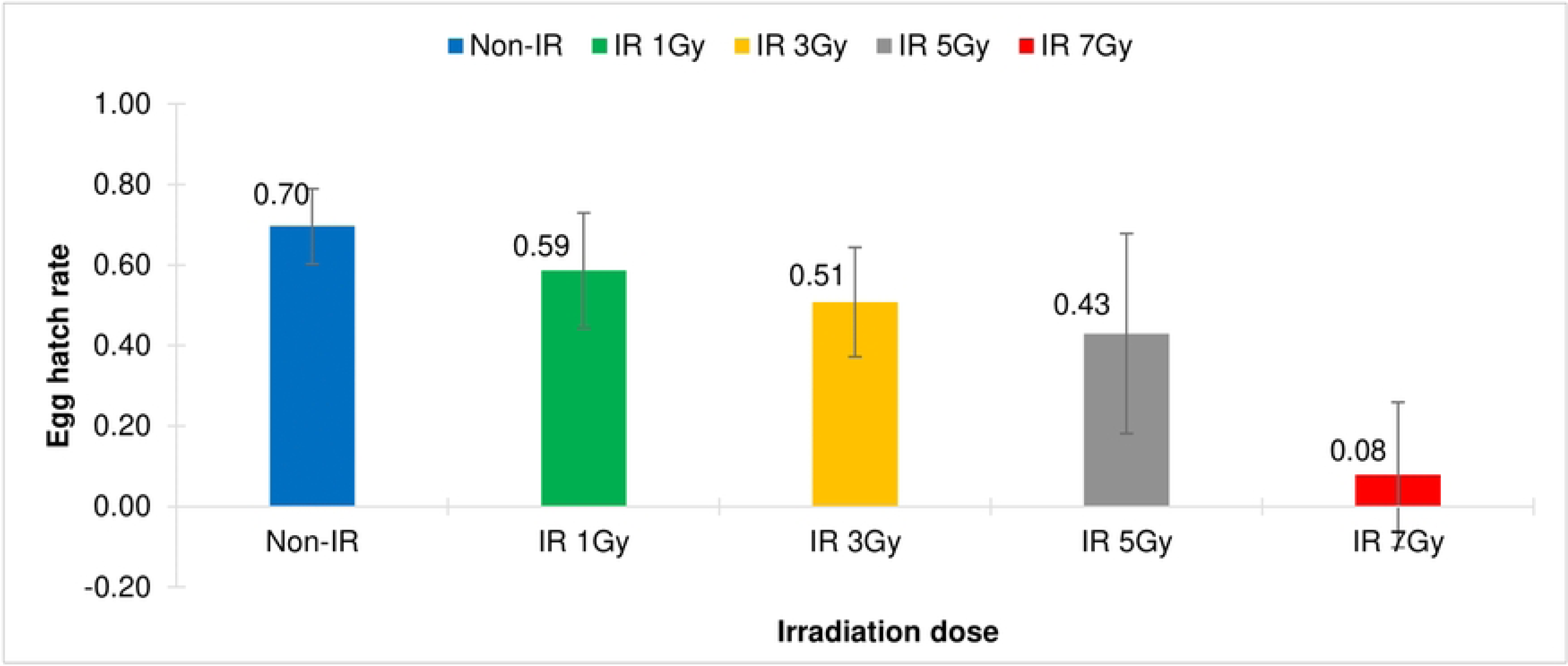
Comparison of egg hatch rates of the cross-mating pairs of the wild-type *Wolbachia* uninfected *Aedes aegypti*, between males emerged from irradiated eggs and non-irradiated females (IR Non-WolB ♂ x Non-IR Non-WolB ♀). Eggs were irradiated with X-ray at the irradiation doses of 1 Gy, 3 Gy, 5 Gy and 7 Gy.

**Table 2.** Comparison of total eggs, total hatched eggs, egg hatch rates and induced sterility of the cross-mating pairs of the wild-type *Wolbachia* uninfected *Aedes aegypti*, between <u>males</u> emerged from irradiated eggs and non-irradiated females (IR Non-WolB ♂ x Non-IR Non-WolB ♀). Eggs were irradiated with X-ray at the irradiation doses of 1 Gy, 3 Gy, 5 Gy and 7 Gy.

| Variables | N | Total eggs<br>(Mean ± SD) | Total hatched eggs<br>(Mean ± SD) | Egg hatch rate<br>(Mean ± SD) | Induce Sterility (IS)<br>(95%CI) |
| --- | --- | --- | --- | --- | --- |
| Non-IR Non-WolB ♂ | 120 | 3,384<br>(112.80 ± 11.87) | 2,349<br>(78.30 ± 11.62) | 0.70<br>(0.70 ± 0.09) | 99.30<br>(-0.73 - -0.66)* |
| IR Non-WolB ♂ 1 Gy | 120 | 3,047<br>(101.57 ± 33.86) | 1,720<br>(57.33 ± 20.62) | 0.59<br>(0.59 ± 0.14)* | 99.41<br>(-0.64 - -0.53)* |
| IR Non-WolB ♂ 3 Gy | 120 | 2,540<br>(84.67 ± 28.51)* | 1,310<br>(43.67 ± 20.34)* | 0.51<br>(0.51 ± 0.14)* | 99.49<br>(-0.56 - -0.46)* |
| IR Non-WolB ♂ 5 Gy | 120 | 2,192<br>(73.07 ± 47.79)* | 1,155<br>(38.50 ± 27.61)* | 0.43<br>(0.43 ± 0.25)* | 99.57<br>(-0.52 - -0.34)* |
| IR Non-WolB ♂ 7 Gy | 120 | 283<br>(9.43 ± 24.59)* | 131<br>(4.37 ± 11.93)* | 0.08<br>(0.08 ± 0.18)* | 99.92<br>(-0.15 - -0.01)* |

\* Significant difference at $p < 0.05$

For sterility of females emerged from irradiated eggs, it was found that the irradiation dose of 7 Gy significantly reduced the total number of eggs of the cross-mating pairs between females emerged from irradiated eggs and non-irradiated males (IR Non-WolB F x Non-IR Non-WolB M), and a significant reduction of egg hatch rates was observed at the irradiation doses starting from 3 Gy onward (0.58 ± 0.10 vs 0.64 ± 1.01, df = 29, t = 2.376, *p*-value = 0.024) (Fig. 3, Table 3); and females became nearly complete sterile at the irradiation dose of 7 Gy (IS = 99.95; 0.05 ± 0.14 vs 0.64 ± 1.01, df = 29, t = 17.124, *p*-value = 0.000).

**Figure 3.**
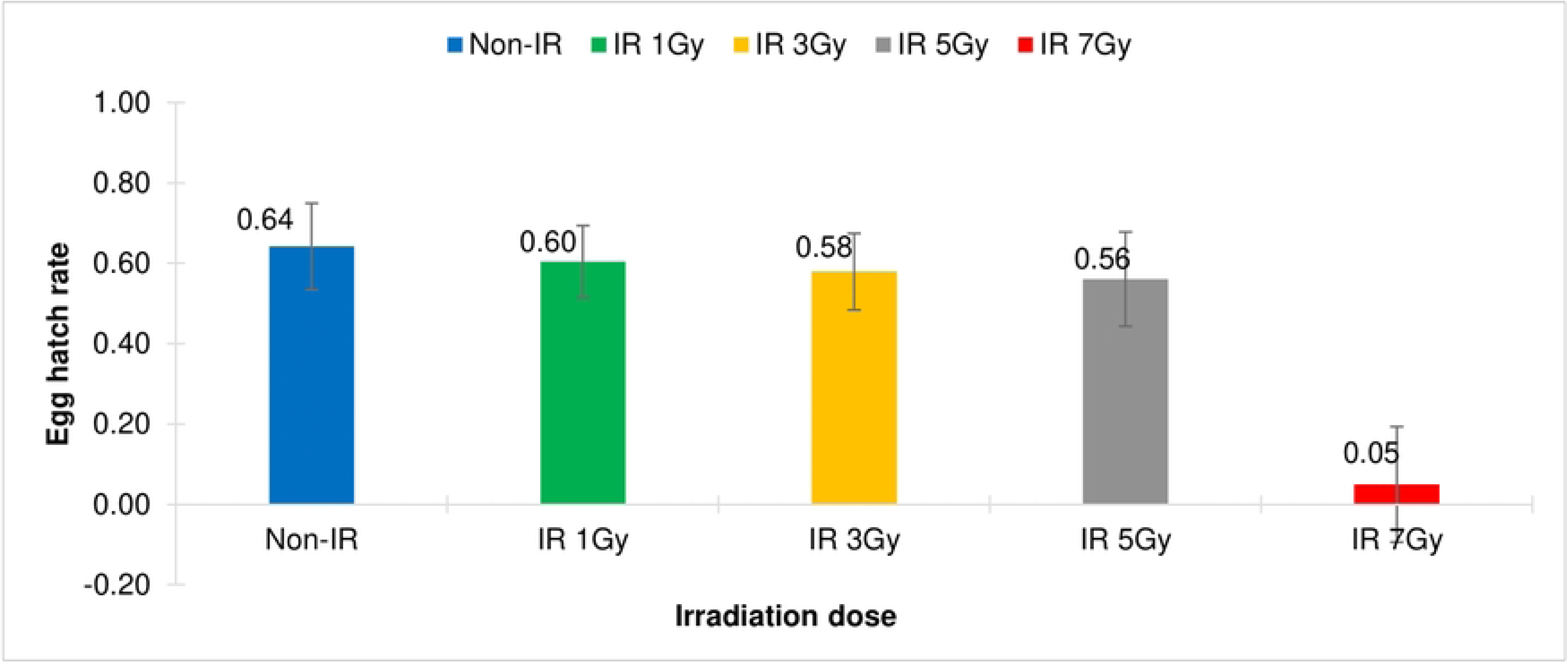
Comparison of egg hatch rates of the cross-mating pairs of the wild-type *Wolbachia* uninfected *Aedes aegypti*, between <u>females</u> emerged from irradiated eggs and non-irradiated males (IR Non-WolB ♀ x Non-IR Non-WolB ♂). Eggs were irradiated with X-ray at the irradiation doses of 1 Gy, 3 Gy, 5 Gy and 7 Gy.

**Table 3.**
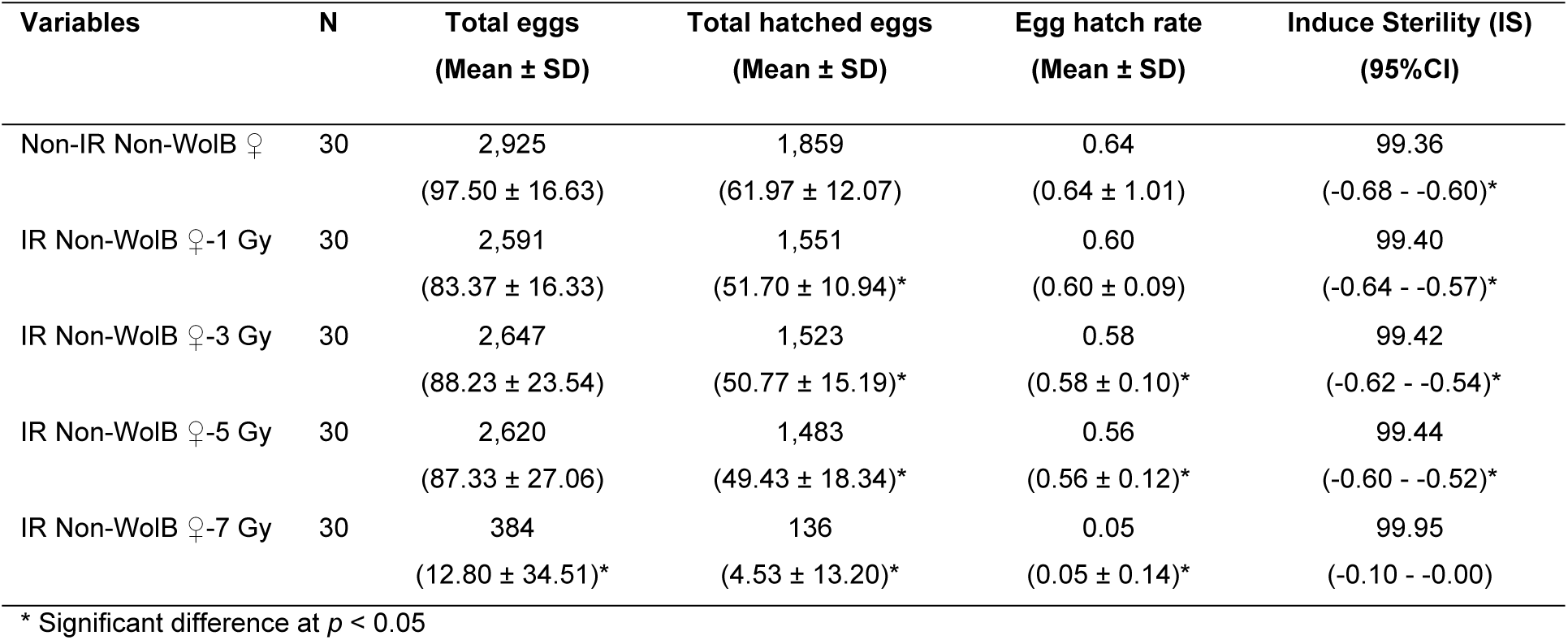
Comparison of total eggs, total hatched eggs, egg hatch rates and induced sterility of the cross-mating pairs of the wild-type *Wolbachia* uninfected *Aedes aegypti*, between <u>females</u> emerged from irradiated eggs and non-irradiated males (IR Non-WolB ♀ x Non-IR Non-WolB ♂). Eggs were irradiated with X-ray at the irradiation doses of 1 Gy, 3 Gy, 5 Gy and 7 Gy.

To summarize, when the wild-type *Wolbachia* uninfected eggs of *Ae. aegypti* were irradiated with X-ray, the irradiation dose of 7 Gy significantly induced more than 92% sterility of both males and females. Therefore, the irradiation dose of 7 Gy could be an optimum dose for irradiation of the wild-type *Wolbachia* uninfected *Ae. aegypti* eggs.

### Development of *Wolbachia* trans-infected *Aedes aegypti* after being irradiated at the egg stage

For an experiment on irradiation of *Wolbachia* trans-infected *Ae. aegypti* eggs, it was found that the irradiation dose from 1 Gy significantly increased the development of the forth-instar larvae when compared to those of the controls (Fig. 4, Table 4). When larvae developed into pupae, significantly high mortality of both male and female pupae was observed at the irradiation dose of 7 Gy when compared to those of the controls.

**Figure 4.**
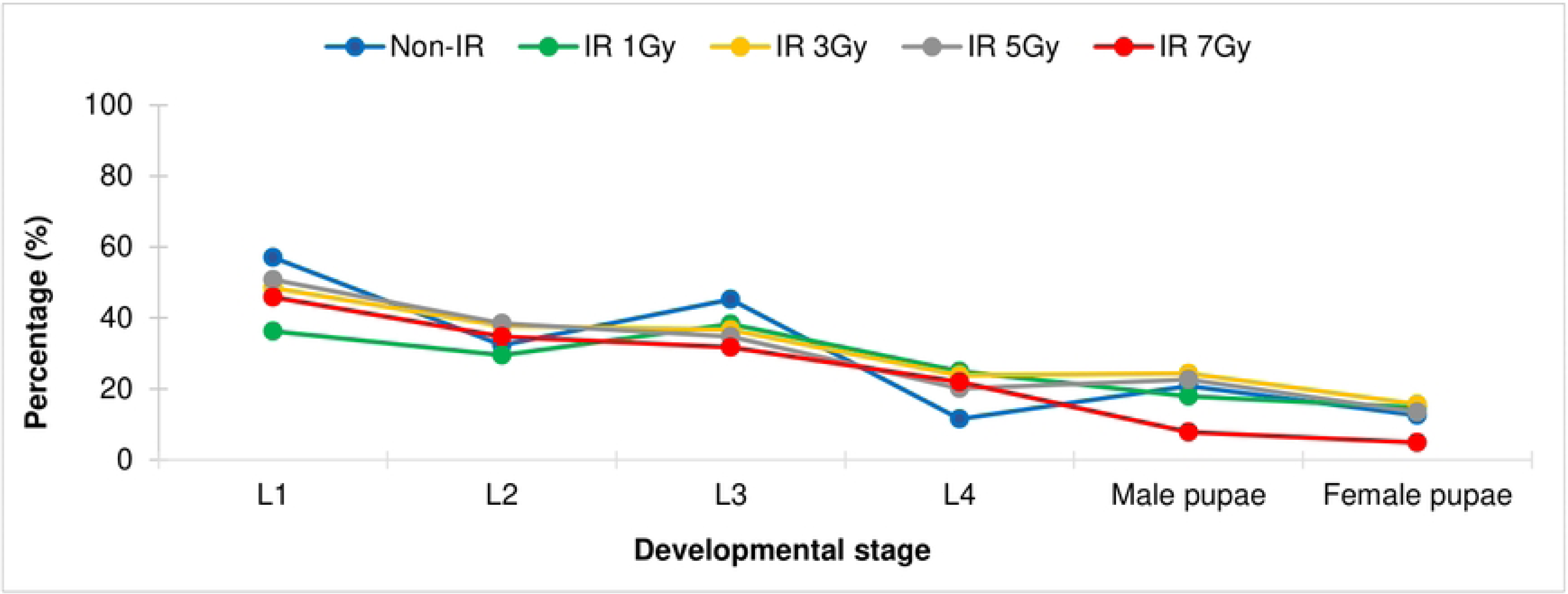
Comparison of larval and pupal development of *Wolbachia* trans-infected *Aedes aegypti* after eggs being irradiated with X-ray at the irradiation doses of 1 Gy, 3 Gy, 5 Gy and 7 Gy.

**Table 4.**
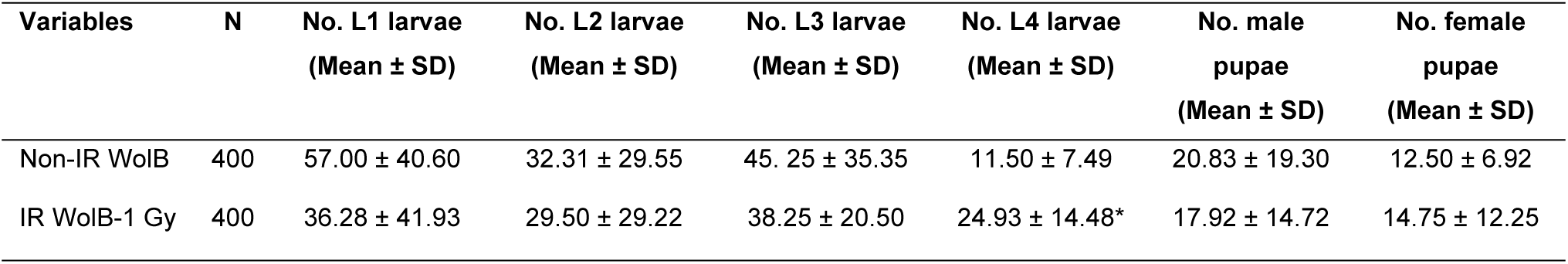
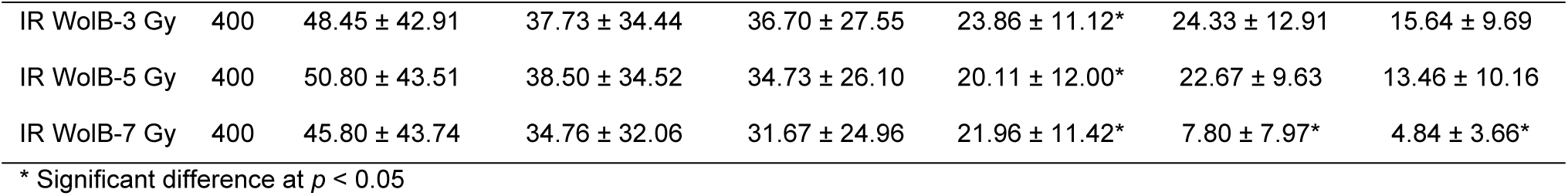
Comparison of larval and pupal development of *Wolbachia* trans-infected *Aedes aegypti* after eggs being irradiated with X-ray at the irradiation doses of 1 Gy, 3 Gy, 5 Gy and 7 Gy.

| Variables | N | No. L1 larvae<br>(Mean ± SD) | No. L2 larvae<br>(Mean ± SD) | No. L3 larvae<br>(Mean ± SD) | No. L4 larvae<br>(Mean ± SD) | No. male pupae<br>(Mean ± SD) | No. female pupae<br>(Mean ± SD) |
| --- | --- | --- | --- | --- | --- | --- | --- |
| Non-IR WolB | 400 | 57.00 ± 40.60 | 32.31 ± 29.55 | 45.25 ± 35.35 | 11.50 ± 7.49 | 20.83 ± 19.30 | 12.50 ± 6.92 |
| IR WolB-1 Gy | 400 | 36.28 ± 41.93 | 29.50 ± 29.22 | 38.25 ± 20.50 | 24.93 ± 14.48* | 17.92 ± 14.72 | 14.75 ± 12.25 |
| IR WolB-3 Gy | 400 | 48.45 ± 42.91 | 37.73 ± 34.44 | 36.70 ± 27.55 | 23.86 ± 11.12* | 24.33 ± 12.91 | 15.64 ± 9.69 |
| IR WolB-5 Gy | 400 | 50.80 ± 43.51 | 38.50 ± 34.52 | 34.73 ± 26.10 | 20.11 ± 12.00* | 22.67 ± 9.63 | 13.46 ± 10.16 |
| IR WolB-7 Gy | 400 | 45.80 ± 43.74 | 34.76 ± 32.06 | 31.67 ± 24.96 | 21.96 ± 11.42* | 7.80 ± 7.97* | 4.84 ± 3.66* |

\* Significant difference at $p < 0.05$

### Sterility of *Wolbachia* trans-infected *Aedes aegypti* after being irradiated at the egg stage

In terms of sterility of males emerged from irradiated eggs, it was found that the irradiation dose of 1 Gy significantly reduced the total number of eggs and the total number of hatched eggs of the cross-mating pairs between males emerged from irradiated eggs and non-irradiated females (IR WolB M x Non-IR WolB F); and a significant reduction of the egg hatch rates was observed when the irradiation dose was at 3 Gy (0.60 ± 0.13 vs 0.70 ± 0.11, df = 29, t = 3.113, *p*-value = 0.004) (Fig. 5, Table 5). Dramatically significant reduction of the egg hatch rates was observed when the irradiation doses were at 5 Gy (0.14 ± 0.29 vs 0.70 ± 0.11, df = 29, t = 10.691, *p*-value = 0.000) and at 7 Gy (0.05 ±0.19 vs 0.70 ± 0.11, df = 29, t = 16.824, *p*-value = 0.000) (Fig. 5, Table 5).

**Figure 5.**
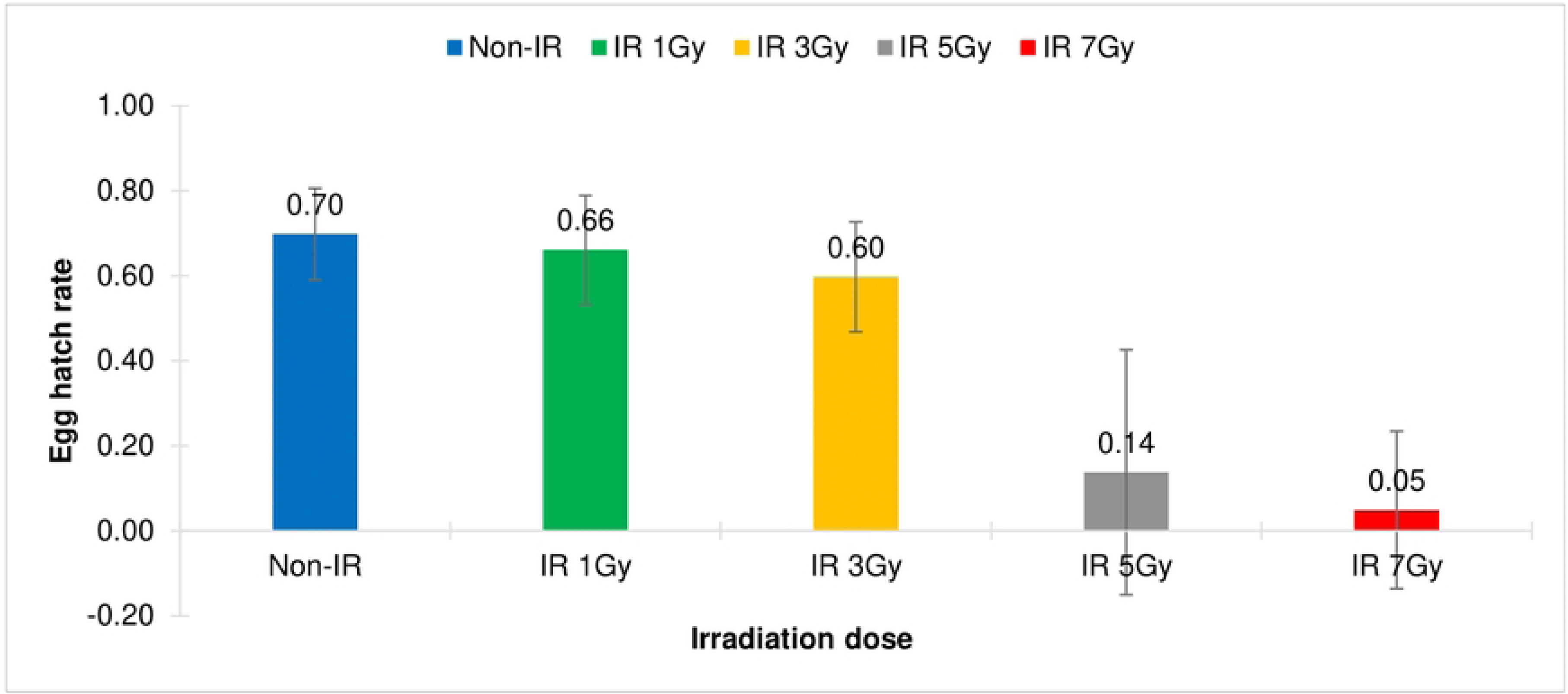
Comparison of egg hatch rates of the cross-mating pairs of *Wolbachia* trans-infected *Aedes aegypti*, between <u>males</u> emerged from irradiated eggs and non-irradiated females (IR WolB ♂ x Non-IR WolB ♀). Eggs were irradiated with X-ray at the irradiation doses of 1 Gy, 3 Gy, 5 Gy and 7 Gy.

**Table 5.**
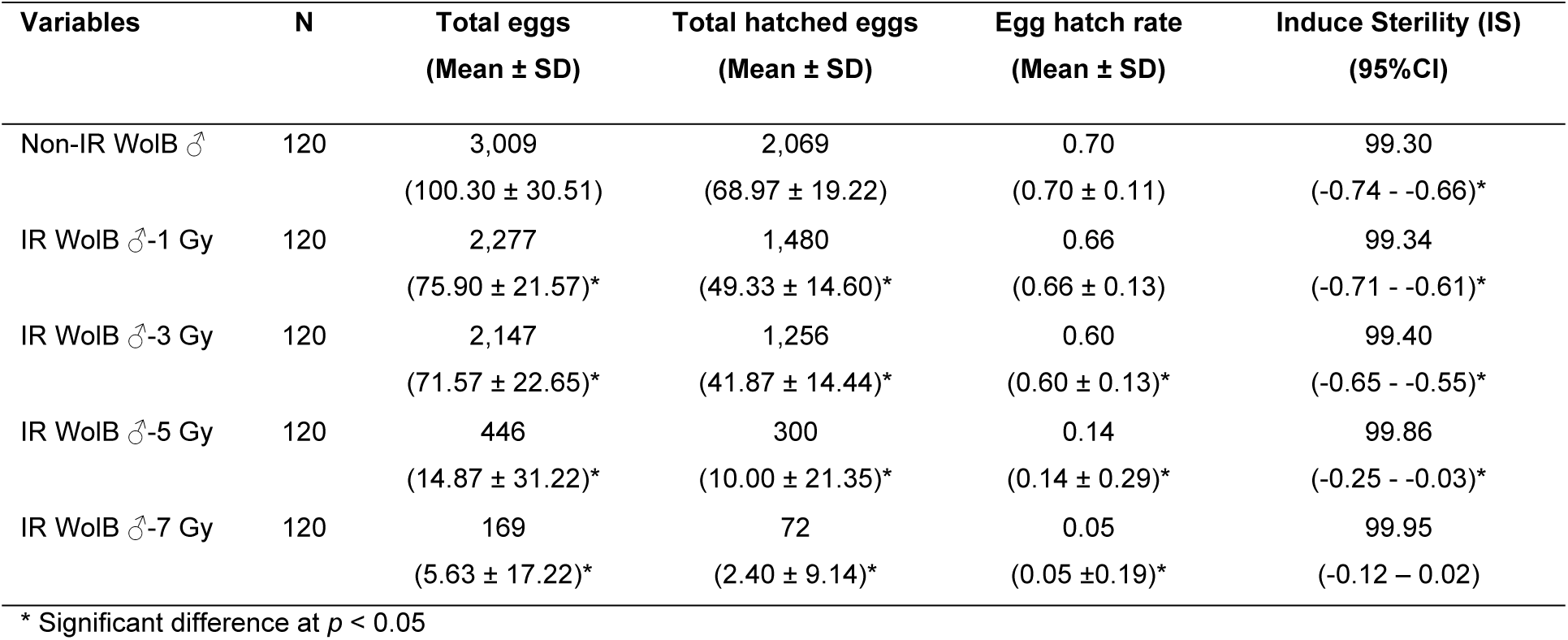
Comparison of total eggs, total hatched eggs, egg hatch rate and induced sterility of the cross-mating pairs of *Wolbachia* trans-infected *Aedes aegypti*, between <u>males</u> emerged from irradiated eggs and non-irradiated females (IR WolB ♂ x Non-IR WolB ♀). Eggs were irradiated with X-ray at the irradiation doses of 1 Gy, 3 Gy, 5 Gy and 7 Gy.

| Variables | N | Total eggs<br>(Mean ± SD) | Total hatched eggs<br>(Mean ± SD) | Egg hatch rate<br>(Mean ± SD) | Induce Sterility (IS)<br>(95%CI) |
| --- | --- | --- | --- | --- | --- |
| Non-IR WolB ♂ | 120 | 3,009<br>(100.30 ± 30.51) | 2,069<br>(68.97 ± 19.22) | 0.70<br>(0.70 ± 0.11) | 99.30<br>(-0.74 - -0.66)* |
| IR WolB ♂-1 Gy | 120 | 2,277<br>(75.90 ± 21.57)* | 1,480<br>(49.33 ± 14.60)* | 0.66<br>(0.66 ± 0.13) | 99.34<br>(-0.71 - -0.61)* |
| IR WolB ♂-3 Gy | 120 | 2,147<br>(71.57 ± 22.65)* | 1,256<br>(41.87 ± 14.44)* | 0.60<br>(0.60 ± 0.13)* | 99.40<br>(-0.65 - -0.55)* |
| IR WolB ♂-5 Gy | 120 | 446<br>(14.87 ± 31.22)* | 300<br>(10.00 ± 21.35)* | 0.14<br>(0.14 ± 0.29)* | 99.86<br>(-0.25 - -0.03)* |
| IR WolB ♂-7 Gy | 120 | 169<br>(5.63 ± 17.22)* | 72<br>(2.40 ± 9.14)* | 0.05<br>(0.05 ± 0.19)* | 99.95<br>(-0.12 - 0.02) |

\* Significant difference at $p < 0.05$

For sterility of females emerged from irradiated eggs, it was found that the irradiation dose of 1 Gy significantly reduced the total number of eggs and the total number of hatched eggs of the cross-mating pairs between females emerged from irradiated eggs and non-irradiated males (IR WolB F x Non-IR WolB M). A significant reduction of the egg hatch rates was observed when the irradiation doses were at 1 Gy (0.45 ± 0.22 vs 0.63 ± 0.11, df = 29, t = 3.940, *p*-value = 0.000) and at 3 Gy (0.16 ± 0.26 vs 0.63 ± 0.11, df = 29, t = 10.020, *p*-value = 0.000) (Fig. 6, Table 6). Moreover, dramatically significant reduction of the egg hatch rates was observed or females became nearly complete sterile when the irradiation dose was at 5 Gy (IS = 99.91, 0.09 ± 0.18 vs 0.63 ± 0.11, df = 29, t = 14.707, *p*-value = 0.000) (Fig. 6, Table 6).

**Figure 6.**
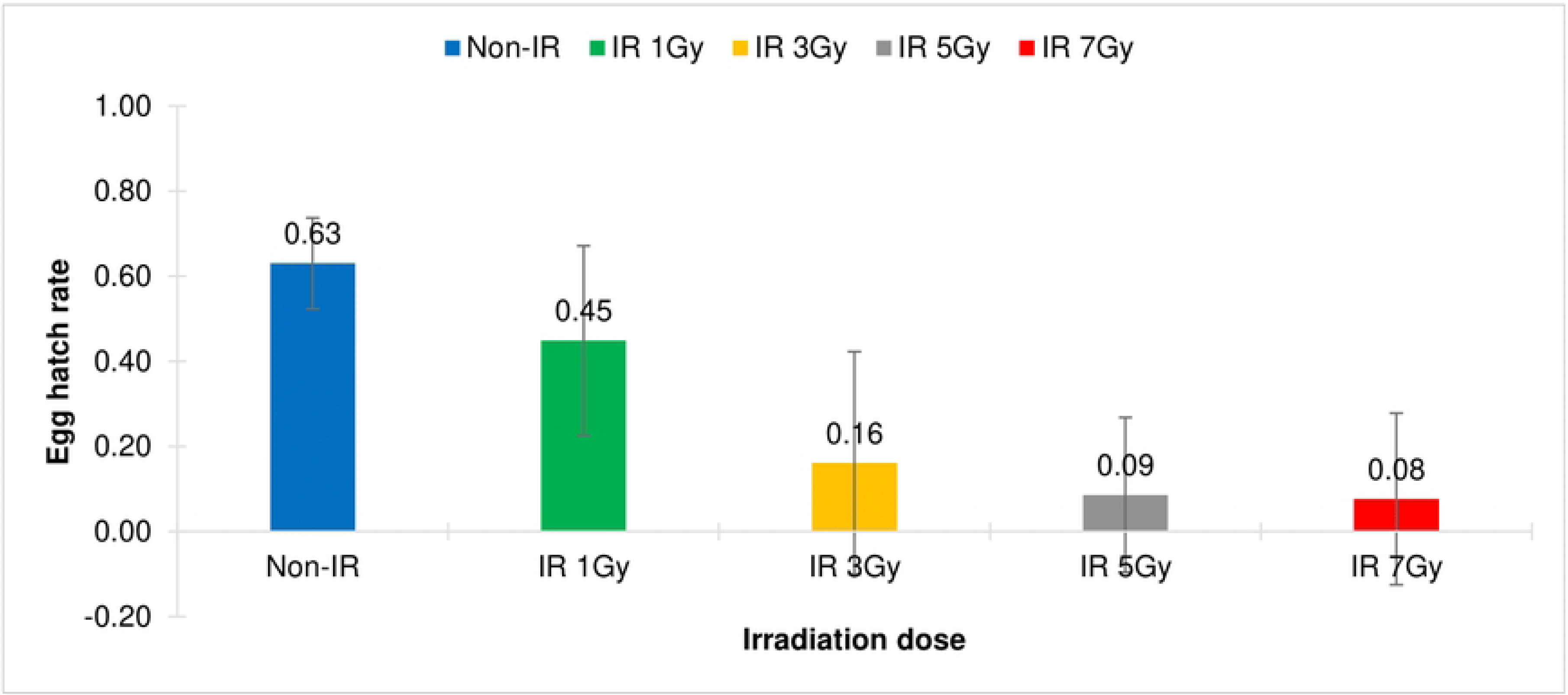
Comparison of egg hatch rates of the cross-mating pairs of *Wolbachia* trans-infected *Aedes aegypti*, between <u>females</u> emerged from irradiated eggs and non-irradiated males (IR WolB ♀ x Non-IR WolB ♂). Eggs were irradiated with X-ray at the irradiation doses of 1 Gy, 3 Gy, 5 Gy and 7 Gy.

**Table 6.** Comparison of total eggs, total hatched eggs, egg hatch rates and induced sterility of the cross-mating pairs of *Wolbachia* trans-infected *Aedes aegypti*, between <u>females</u> emerged from irradiated eggs and non-irradiated males (IR WolB ♀ x Non-IR WolB ♂). Eggs were irradiated with X-ray at the irradiation doses of 1 Gy, 3 Gy, 5 Gy and 7 Gy.

| Variables | N | Total eggs<br>(Mean ± SD) | Total hatched eggs<br>(Mean ± SD) | Egg hatch rate<br>(Mean ± SD) | Induced Sterility<br>(IS)<br>(95%CI) |
| --- | --- | --- | --- | --- | --- |
| Non-IR WolB ♀ | 120 | 3,130<br>(104.33 ± 27.58) | 1,944<br>(64.80 ± 16.43) | 0.63<br>(0.63 ± 0.11) | 99.37<br>(-0.67 - -0.59)* |
| IR WolB ♀-1 Gy | 120 | 2,371<br>(79.03 ± 25.53)* | 1,080<br>(36.00 ± 22.42)* | 0.45<br>(0.45 ± 0.22)* | 99.55<br>(-0.53 - -0.37)* |
| IR WolB ♀-3 Gy | 120 | 1,997<br>(66.57 ± 38.91)* | 419<br>(13.97 ± 23.53)* | 0.16<br>(0.16 ± 0.26)* | 99.84<br>(-0.26 - -0.06)* |
| IR WolB ♀-5 Gy | 120 | 1,540<br>(51.33 ± 46.79)* | 173<br>(5.77 ± 12.04)* | 0.09<br>(0.09 ± 0.18)* | 99.91<br>(-1.54 - -0.02)* |
| IR WolB ♀-7 Gy | 120 | 361<br>(12.03 ± 25.35)* | 141<br>(4.70 ± 12.88)* | 0.08<br>(0.08 ± 0.20)* | 99.92<br>(-0.15 - -0.00)* |

\* Significant difference at $p < 0.05$

In summary, the X-ray irradiation dose of 7 Gy significantly induced male sterility, whereas the irradiation dose of only 5 Gy significantly induced female sterility. Therefore, the X-ray irradiation dose of 5 Gy could be an optimum dose for irradiation of *Wolbachia* trans-infected *Ae. aegypti* eggs as it could induce high level of sterility in males and nearly complete sterility in females.

## Discussion

Irradiation on different mosquito species at various stages of life cycle showed increased or declined effect on adult life span including its subsequent generations [23, 35]. Age had been shown to have an effect on radio-sensitivity, and the older the life stage, or the age within the life stages, the more radio-resistant the insects became [18]. Treatment of early life stages of mosquitoes and environmental changes greatly influenced lifespan of adult mosquitoes [20], so in order to reduce somatic damage, irradiation of insects at or near to the completion of their development, i.e., the late pupal and adult stages of mosquitoes, had been emphasized [19, 20, 24, 28, 36]. However, the optimum developmental stage for irradiation depended on many factors including ease of handling on a mass-production scale, logistics of the irradiation process (e.g., the need to irradiate large numbers of insects), competitiveness of the insect, release methodology, and costs [24]. In this study, we investigated the effect of irradiation at the egg stage on sterility of *Wolbachia* trans-infected and uninfected *Ae. aegypti* in order to find an appropriate and more convenient methods for mass production of sterile males prior to the release of sterile males for vector control.

So far very few information has been available for irradiation of eggs, especially for mosquitoes. Due to inefficient results in controlling *Aedes*-transmitted arboviral diseases, research on irradiation of mosquito stages beside adults and immatures was needed [37]. In this study, we observed a late pupation and high mortality in pupae when *Ae. aegypti* eggs were irradiated with X-ray at the irradiation dose of 7 Gy. Our results were supported by Tantawy et al. [38] who showed unacceptable mortality when eggs of *Anopheles pharoensis* were irradiated with gamma radiation [35]. Moreover, our studies showed that the irradiation dose of 7 Gy significantly induced more than 92% sterility in the wild-type *Wolbachia* uninfected *Ae. aegypti* males and females after being irradiated at the egg stage. In terms of *Wolbachia* trans-infected *Ae. aegypti*, the same irradiation dose of 7 Gy also induced sterility in males and lower dose of 5 Gy was sufficient to induce complete sterility in females. Our results were supported by the study of Akter and Khan [25] who found that low dose radiation from 1-10 Gy had significant effect on pupation and adult emergence when they were irradiated at eggs. Our results coincided with the study of Furaki et al. [39] who observed a reduction in egg hatching as well as in adult emergence of the flour beetles and the almond moth when their eggs were exposed to UV irradiation. The same study concluded that damaging of the surface tissues of the eggs by radiation could be fatal, especially at the advanced stages of development; as development proceeded the embryonic regions became more specialized, so different organ fields could no longer replace each other [39].

Although pupae are the preferred stages to sterilize during SIT operations, but due to limited pupation time, the distances for their distribution were limited to closer localities where the irradiation source was located [37]. In contrary, eggs could be distributed to logistically remote field sites where they can hatch, develop into adults, and fly out to compete with wild mosquitoes [40]. Moreover, the diapause of mosquito eggs was considered an important factor hindering mosquito population suppression [41]. The cost-benefit of using the egg stage in field distribution was very promising to low-income endemic countries. Hence, the expansion of sterilized insect programs could be facilitated by using either irradiated eggs or *Wolbachia* vertical infection [37]. Our results showed an effect of low-dose X-ray irradiation on sterility of both male and female wild-type and *Wolbachia* trans-infected *Ae. aegypti* when eggs were irradiated, and this approach could be applied for SIT operation for vector control in the fields.

## Acknowledgments

Experiments were reviewed and approved by the Faculty of Science, Mahidol University-Institutional Animal Care and Use Committee (MUSC-IACUC) (MUSC64-044-593). The authors would like to thank Ms. Natchaya Klinpikul and Mr. Kuang Chalongpak for mosquito rearing, Ms. Kanyarat Yimpramote and Ms. Kamolrat Tharaporn for lab assistance, and Mr. Thodsapon Thannarin for irradiation process.

